# Estimating somatic mutation rates by Duplex Sequencing in non-model organisms: *Daphnia magna* as a case study

**DOI:** 10.1101/2022.05.31.494242

**Authors:** Eli Sobel, Jeremy E. Coate, Sarah Schaack

**Author notes:** **Corresponding author:** Sarah Schaack, Reed College, Department of Biology, 3203 SE Woodstock Blvd, Portland, OR, 97202, USA.

## Abstract

Somatic mutations are evolutionarily important as determinants of individual organismal fitness, as well as being a focus of clinical research on age-related disease, such as cancer. Identifying somatic mutations and quantifying mutation rates, however, is extremely challenging and genome-wide somatic mutation rates have only been reported for a few model organisms. Here, we describe the application of Duplex Sequencing on bottlenecked WGS libraries to quantify genome-wide somatic base substitution rates in *Daphnia magna. Daphnia*, historically an ecological model system, has more recently been the focus of mutation studies, in part because of its high germline mutation rates. Using our protocol and pipeline, we estimate a somatic mutation rate of 2.14 × 10^−7^ substitutions per site (in a genotype where the germline rate is 3.60 × 10^−9^ substitutions per site per generation). To obtain this estimate, we tested multiple dilution levels to maximize sequencing efficiency, and developed bioinformatic filters needed to minimize false positives when a high quality reference genome is not available. In addition to laying the groundwork for estimating genotypic variation in rates of somatic mutations within *D. magna*, we provide a framework for quantifying somatic mutations in other non-model systems, and also highlight recent innovations to single molecule sequencing that will help to further refine such estimates.

## Introduction

Efficient methods for detecting rare genetic variants are critical for both clinical applications and for basic biology [1,2]. Next generation sequencing (NGS) has been used extensively to identify germline variants, but the variant allele fraction (VAF) of many somatic mutations (in some cases <0.01%) is well below the error rates associated with standard NGS, making the detection of somatic mutations challenging [1,3–5]. Single-molecule sequencing (SMS) technologies, however, dramatically reduce the error rates associated with high-throughput sequencing, potentially enabling the interrogation of rare, subclonal variation by NGS (e.g., Safe-SeqS [1], Duplex Sequencing [6,7], smMIP [2], BotSeqS [8], Hawk-Seq [9], PECC-Seq [10], and NanoSeq [11]).

Generally, SMS methods reduce error rates by uniquely barcoding individual DNA molecules. Amplification of these uniquely barcoded templates produces PCR duplicates that, upon sequencing, can be grouped into ‘read families’ based on their shared barcodes (Unique Molecular Identifiers [UMIs]) [12]. Mutations present in the original molecule should be present in the majority of PCR duplicates, whereas errors introduced by PCR and sequencing will typically only be observed in a small subset. Most artifacts, therefore, can be identified and removed in the process of constructing a consensus sequence from each read family [1]. UMIs can be endogenous (e.g., the random sites of fragmentation at each end of a DNA molecule generated during library preparation) or exogenous (random ‘barcode’ sequences incorporated during library construction).

The accuracy of SMS approaches is further enhanced by Duplex Sequencing [6,7,13,14], wherein the two strands of a target molecule are labeled with complementary UMIs so that read families, and consensus sequences, can be generated separately for each strand of the original template (Single Strand Consensus Sequences, SSCSs; **Fig. 1**). Complementary SSCSs can then be compared to generate Duplex Consensus Sequences (DCSs). Because true mutations alter the sequence of both strands, only complementary variants present in both read families are scored as mutations. In contrast to the first generation of SMS, Duplex Sequencing enables detection of errors that arise even in the first round of PCR. Schmitt et al. [6] showed that although non-duplex-based SMS eliminated >99% of technical errors, 90% of the remaining mutations were still artifacts. Applying Duplex Sequencing, however, further eliminated ∼90% of mutations identified in single-strand consensus sequences. The theoretical background error rate of the duplex approach is <1 error per 10^9^ nucleotides. Thus, first generation SMS methods, and, to a greater extent, Duplex Sequencing, filter out the vast majority of artifacts, while effectively detecting variants with low VAF [1,2,6].

**Figure 1.**
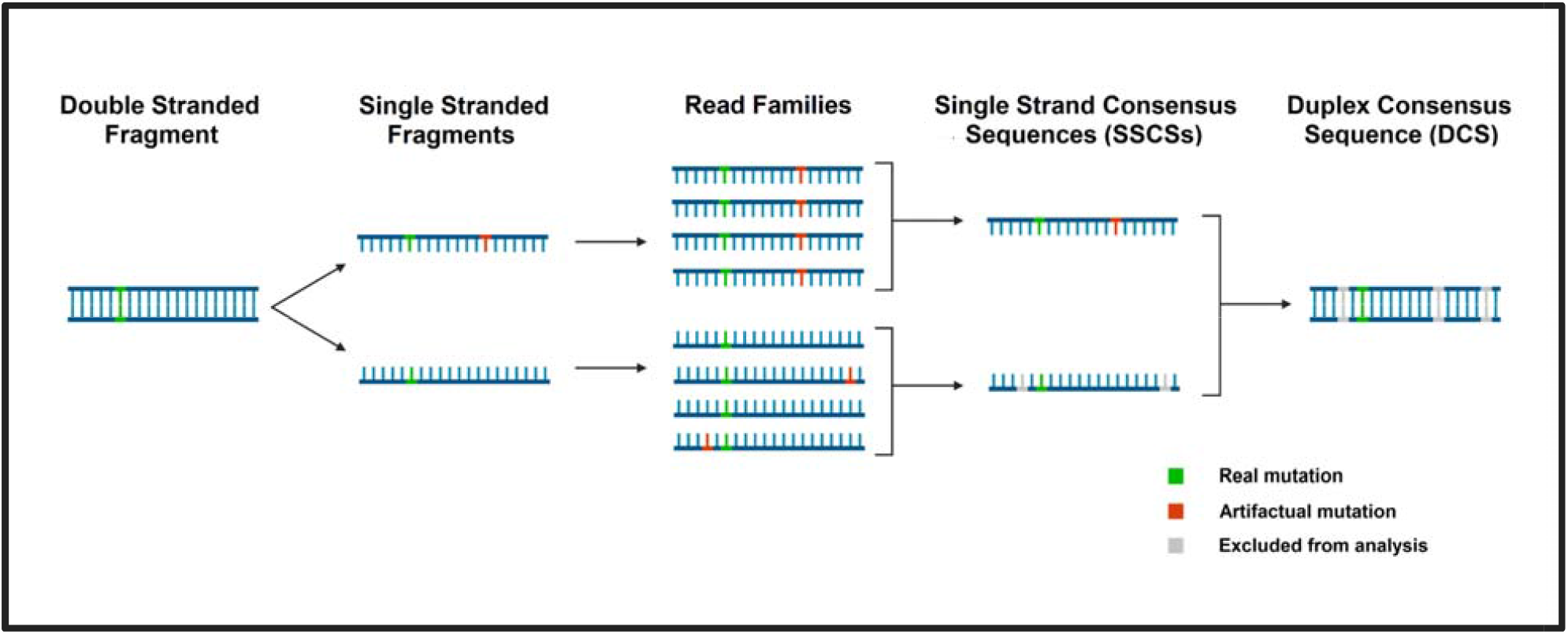
The Duplex Sequencing consensus-making process, where consensus sequences are formed from read families. Both strands of a DNA fragment are amplified and the resulting amplicons are compared to build a single strand consensus sequence (SSCS). The SSCSs generated from the two strands of the original fragment are then compared to one another to build a duplex consensus sequence (DCS). All reads must agree to form a consensus base for any given position. Blue lines represent strands of genomic DNA (gDNA), green notches represent bona fide mutations, red notches represent artifacts (PCR or sequencing errors), and gray notches indicate bases that have been marked as N and excluded from analysis.

However, two significant challenges remain when using Duplex Sequencing to estimate genome-wide somatic mutation rates. First, because SMS relies on sequencing of multiple PCR amplicons per template, it becomes prohibitively expensive as a method for comprehensively surveying large genomes [2,7]. In fact, many of these methods were specifically designed for assaying small genomes (<1-2 Mbp; [7,13]) or small target regions of large genomes [1,2,7,15]. Diluting (“bottlenecking”) samples prior to library construction is an additional refinement that allows for unbiased sub-sampling of the genome [8,9], reducing the amount of sequencing required to infer the genome-wide mutation rate (**Fig. 2**). In bottleneck approaches, DNA is diluted to a very low (attomolar) concentration. Drastically reducing the number of molecules in the library prior to amplification increases the fraction of remaining molecules that are sequenced redundantly (more than one PCR duplicate is sequenced per original DNA fragment). Dilution also produces a more even distribution of read family sizes [8]. Thus, bottlenecking is a simple way to reduce the ratio of raw reads to read families, thereby optimizing sequencing efficiency (the number of instructive read families generated per raw read) while sampling the genome in an unbiased fashion. However, determining the optimal dilution, the point at which the number of DCSs per raw read is maximized, is not trivial, and must be determined empirically by evaluating the sequencing efficiencies of libraries with a range of dilution levels.

**Figure 2.**
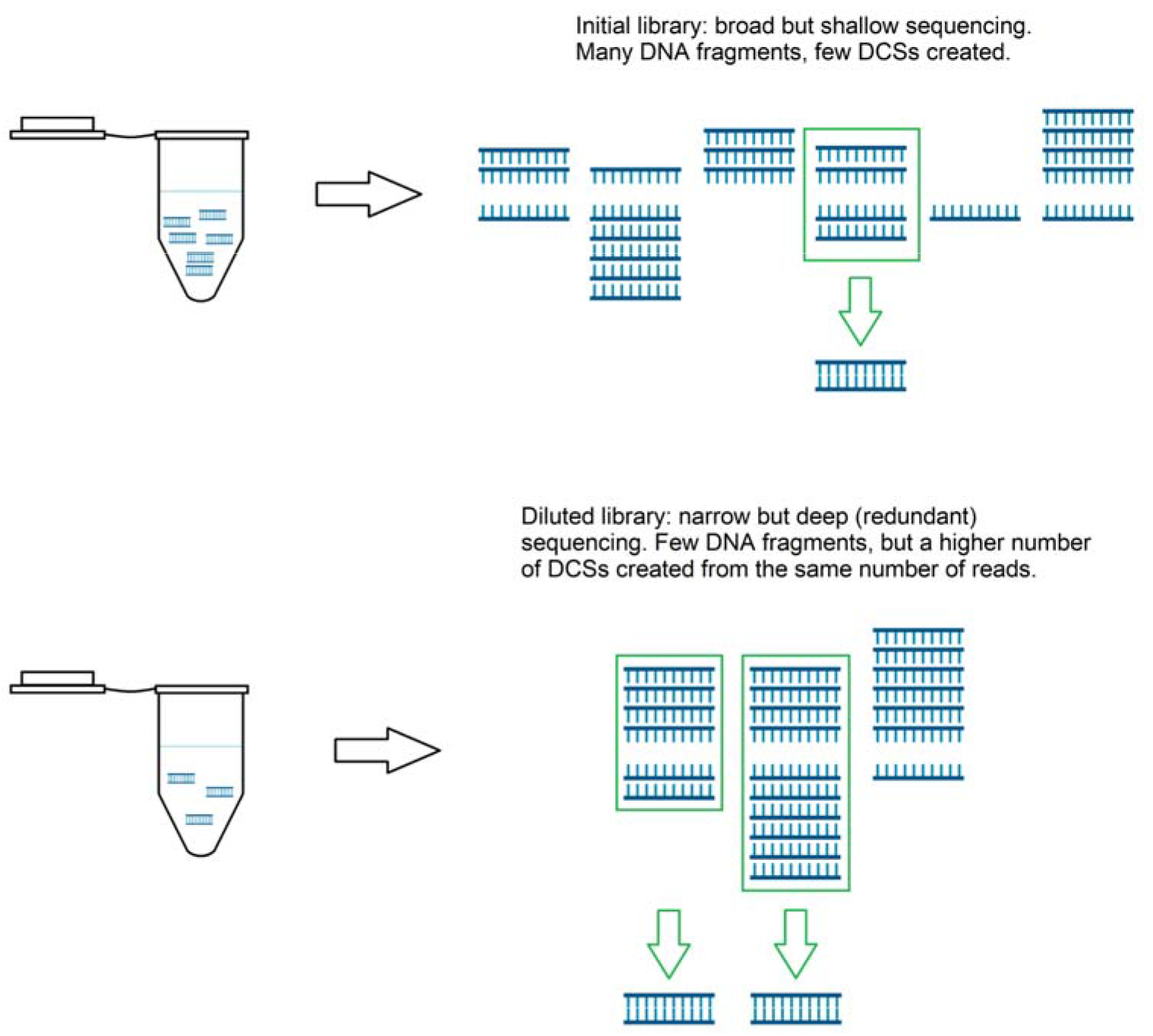
Schematic showing the importance of optimizing dilution for increased efficiency. The distribution of PCR duplicates produced per dsDNA fragment in undiluted (top) versus diluted (bottom) gDNA libraries. Blue double stranded combs in the tube represent dsDNA fragments, while single stranded blue combs represent PCR duplicates produced from the top and bottom strands of those fragments. In the undiluted library, many fragments are not sufficiently amplified and/or sequenced and thus cannot be used to build DCSs. In the diluted library, a high number of PCR duplicates per fragment are produced, allowing more DCSs to be produced (green arrows) from the same number of reads.

Second, in addition to filtering out artifactual mutations resulting from library preparation and sequencing, the inference of mutations is only as robust as the reference genome to which read families are compared. To date, only a few studies have utilized bottlenecked Duplex Sequencing to estimate genome-wide mutation rates, and these studies have been limited to humans and mice [6,8–10]. By virtue of their focus on model organisms, all genome-wide Duplex Sequencing studies have had at their disposal high-quality, chromosome-level reference genomes. For most organisms, however, reference genomes of draft quality have hundreds or thousands of unordered scaffolds, likely containing numerous sequence and assembly errors and gaps. This adds a significant layer of complexity, and source of error, when trying to estimate somatic mutation rates, and strategies for adapting SMS approaches to non-model systems with draft-quality genomes are needed.

Here, we provide guidance to address both of these challenges. First, we provide an example for how to identify the optimal dilution to maximize sequencing efficiency, and discuss how this optimum is influenced by the stringency of parameters used for DCS building. Second, we develop bioinformatic parameters to mask problematic regions of the reference genome and limit our analysis to high-confidence regions in which somatic mutations could be called accurately. In doing so, we describe the application of bottlenecked Duplex Sequencing to estimate a genome-wide somatic mutation rate in *Daphnia magna*, thereby providing a framework for expanding the use of SMS to non-model systems.

## Methods

### Study System and Experimental Design

*Daphnia magna* descended from an individual collected in Israel (genotype ‘IA’; Ho et al. 2020) were reared in a 50 ml conical tube containing 35 ml of Aachener Daphnien Medium (ADaM) [16], and kept in a Percival™ environmental chamber set at 18C and 16:8 hour light:dark in the Schaack Lab (Reed College). Animals were fed algae (*Scenedesmus obliquus*) twice per week. At 60 days old, three individuals with empty brood pouches were collected in a single 1.5 ml microcentrifuge tube, flash frozen in liquid nitrogen and stored at -80°C prior to DNA extraction.

### DNA isolation and quantification

DNA was extracted with DNAzol (Molecular Research Center, Inc.; DN127) as follows: Frozen samples were pulverized with a plastic pestle, then mixed with 200 µl of DNAzol and incubated at room temperature for 10 min. 100 µl of 100% EtOH was added to each sample, mixed thoroughly and centrifuged for 5 minutes at 15,000 x g. Supernatant was discarded, and the pellet was washed with a mixture of 70% DNAzol and 30% EtOH, followed by a second wash with 70% EtOH. After removing the supernatant, the pellet was air dried for 1 minute, then resuspended in 30 µl TE buffer (10mM Tris-HCl, pH 8.0; 0.1mM EDTA). DNA was treated with RNAse I_f_ (New England Biolabs; M0243) for 10 minutes at room temperature, purified using the DNA Clean & Concentrator-25 Kit (Zymo Research; D4033) according to the manufacturer’s protocol, and eluted in 30 µl TE buffer. DNA concentration was measured with a Qubit 4.0 using the Qubit 1x dsDNA High Sensitivity kit (Thermo Fisher Scientific; Q33230).

### Library Construction

The initial steps of library construction, up to PCR amplification, were performed using the NEBNext Ultra II FS DNA Library Prep kit (New England Biolabs; E6177) according to the manufacturer’s protocol, using 115 ng of DNA as input. The DNA was subjected to a 15 min fragmentation period and size selected using the sample purification beads. After fragmentation and adapter ligation, unamplified library concentration was assessed by Qubit as described above, and fragment size distribution was assessed by Agilent 2100 Bioanalyzer and High Sensitivity (HS) DNA Chip at the GPSS core facility at Oregon Health & Sciences University. Based on the concentration estimated by Qubit and average fragment size estimated by Bionanalyzer, the library was serially diluted to 1 fmol/15 µl, 150 amol/15 µl, 100 amol/15 µl, 50 amol/15 µl, and 10 amol/15 µl. 15 µl of each diluted library was then amplified in a 50 µl PCR reaction to append unique dual indices on each DNA fragment and produce multiple copies per fragment. For this step, if too few PCR cycles are used, insufficient template will be generated for sequencing. Conversely, over-amplification has been shown to produce high molecular weight artifacts [7]. Because the optimal number of cycles is dictated by the amount of input DNA, each dilution required a different number of cycles. Based on the guidelines in the NEBNext Ultra II FS library protocol (Dual Index Kit 1; New England Biolabs; NEB#7600S), the 1 fmol, 150 amol, 100 amol, 50 amol, and 10 amol libraries were amplified for 11, 14, 14, 16, and 18 PCR cycles, respectively, which yielded ca. 100 ng of DNA per library while avoiding high MW artifacts. Libraries were assessed by Agilent TapeStation at MedGenome, and average fragment sizes were 430 bp, 415 bp, 387 bp, 423 bp, and 287 bp for the 1 fmol, 150 amol, 100 amol, 50 amol, and 10 amol libraries, respectively. Libraries were pooled and sequenced (2×150 bp paired end) on an Illumina HiSeq X at MedGenome (https://research.medgenome.com/).

### Bioinformatic Analysis

The bioinformatic pipeline to generate consensus reads was developed by Brendan Kohrn and the Kennedy lab [7]; https://github.com/Kennedy-Lab-UW/Duplex-Seq-Pipeline) and was implemented with the following modifications and parameters. The first 12 bases from each read in a read pair (which define the first and last 12 bases of the sequenced insert) were used as endogenous molecular barcodes, in contrast to ligating exogenous barcodes to the ends of inserts during library preparation as described by Kennedy et al. [7]. Additionally, 15 bp were cut from the 5’ ends of reads (including the 12bp used to generate duplex tags) prior to mapping. This additional end clipping is based on empirical observations that mutations are found in excess near the ends of inserts that are likely explained by errors introduced as a result of DNA fragmentation and end repair [7,11]. Reads were then aligned to the reference genome for the IA genotype assembled by Ho et al. [17] using bwa-mem [18,19]. Unmapped reads (bitwise flag values of 77 and 141) were filtered out of the dataset.

Reads that mapped to the same genomic location and shared identical endogenous barcodes were considered PCR duplicates of the same DNA insert and were grouped into read families, with separate families generated for R1 and R2 reads. Within a read family, only reads sharing the most common Compact Idiosyncratic Gapped Alignment Report (CIGAR) string were retained. Read families were discarded if fewer than two such reads remained. For the remaining read families, the retained reads were used to generate a single-strand consensus sequence (SSCS). SSCSs mapping to the same location in the genome and with complementary endogenous barcodes (indicating they were produced from opposite strands of the same insert) were then used to make duplex consensus sequences (DCSs). During both single strand and duplex consensus making, all reads were required to agree at a given position, otherwise that base was marked as N. DCSs were then aligned to the reference genome using bwa-mem [18,19], and any unmapped DCSs were filtered out of the dataset. In addition to filtering out unmapped DCSs, we removed improperly paired alignments, and DCSs were required to have a minimum mapping quality of 55, an insert size between 20 and 500 bp, and to align uniquely to a single locus. Non-unique alignments were filtered out by requiring consensus reads to have a suboptimal alignment score (XS:i flag assigned by bwa-mem) equal to zero. Variant calling was then performed on the consensus reads using bcftools [20,21]. For determining the optimal dilution, a random sample of 50 million read pairs from each library was run through the pipeline as described above.

### Identifying and masking problematic genomic intervals

Variants were only considered to be valid if they fell outside masked regions of the genome. We excluded variants at heterozygous sites identified by Ho et al. [17], and identified genomic intervals for masking in 3 different ways: regions flagged by RepeatModeler or RepeatMasker [22,23], regions of unusually high coverage in the reference genome assembly, and windows of 500 bp containing more than 3 variants. Regions with high coverage in the reference genome were masked, since they represent sites of potential mapping errors. Such regions were identified by mapping the reads used to build the reference assembly back to the reference assembly, generating coverage graphs using bedtools *genomecov*, and masking intervals with more than double the expected coverage of 40x.

To identify regions with a high density of variants, the genome was divided into 500 bp sliding windows, incrementing by 100 bp, using bedtools *makewindows* [24], and intervals with >3 variants were identified. Once bed files of all four intervals to be masked were made, the intervals in each bed file were extended by 500 bp using the bedtools *slop* utility [24]. Then, we used bedtools to merge and sort the three sets of filtered intervals, and subtracted them from the bed file of the whole genome to generate the masked genome. To identify variants in the masked genome, we intersected the masked genome with the list of initial variants called by bcftools. Each of the 120 remaining variants was manually inspected using IGV [25]. We manually excluded variants with a variant allele fraction over 0.5 in cases where a locus was covered by multiple DCSs, or where the reads used to build the reference genome showed heterozygosity at that locus. Variants in regions of the reference assembly not covered by reads used to build the reference assembly were also manually masked.

## Results and Discussion

### Optimizing efficiency

The goal of bottlenecking (diluting) a genomic library is to optimize the number of consensus sequences generated per read (**Fig. 2**). Because Duplex Sequencing relies on redundantly sequencing individual DNA fragments, optimizing the dilution is a critical step in bottleneck sequencing. The optimal amount of input DNA should strike a balance between being low enough that a large fraction of fragments are amplified and sequenced with some redundancy, but high enough to avoid excessive redundancy. When only one copy of a DNA fragment is sequenced it cannot be analyzed because no consensus sequence can be generated. Conversely, over-dilution results in unnecessary and wasteful resequencing of the few DNA fragments retained in the library.

### The lowest DNA input (10 amol) was most efficient

To identify the optimal input of DNA for maximizing sequencing efficiency, we generated a single DNA library then serially diluted it to obtain five different DNA input amounts for PCR amplification (1 fmol, 150 amol, 100 amol, 50 amol, and 10 amol) and sequencing. Consensus-making efficiency varies with the number of input reads [8,9], so it is necessary to control for differences in input read number when evaluating the sequencing efficiencies of different dilutions. Because the number of read pairs produced per library varied from 51.7-115.5 M, we downsampled four of the five libraries (excluding the 150 amol library) to 50M read pairs prior to consensus building and comparison (**Table 1**). We did not analyze a downsampled 150 amol library because a clear picture emerged from the other four libraries (**Fig. 3**) that an input of 10 amol or less is optimal (see below).

**Table 1.**
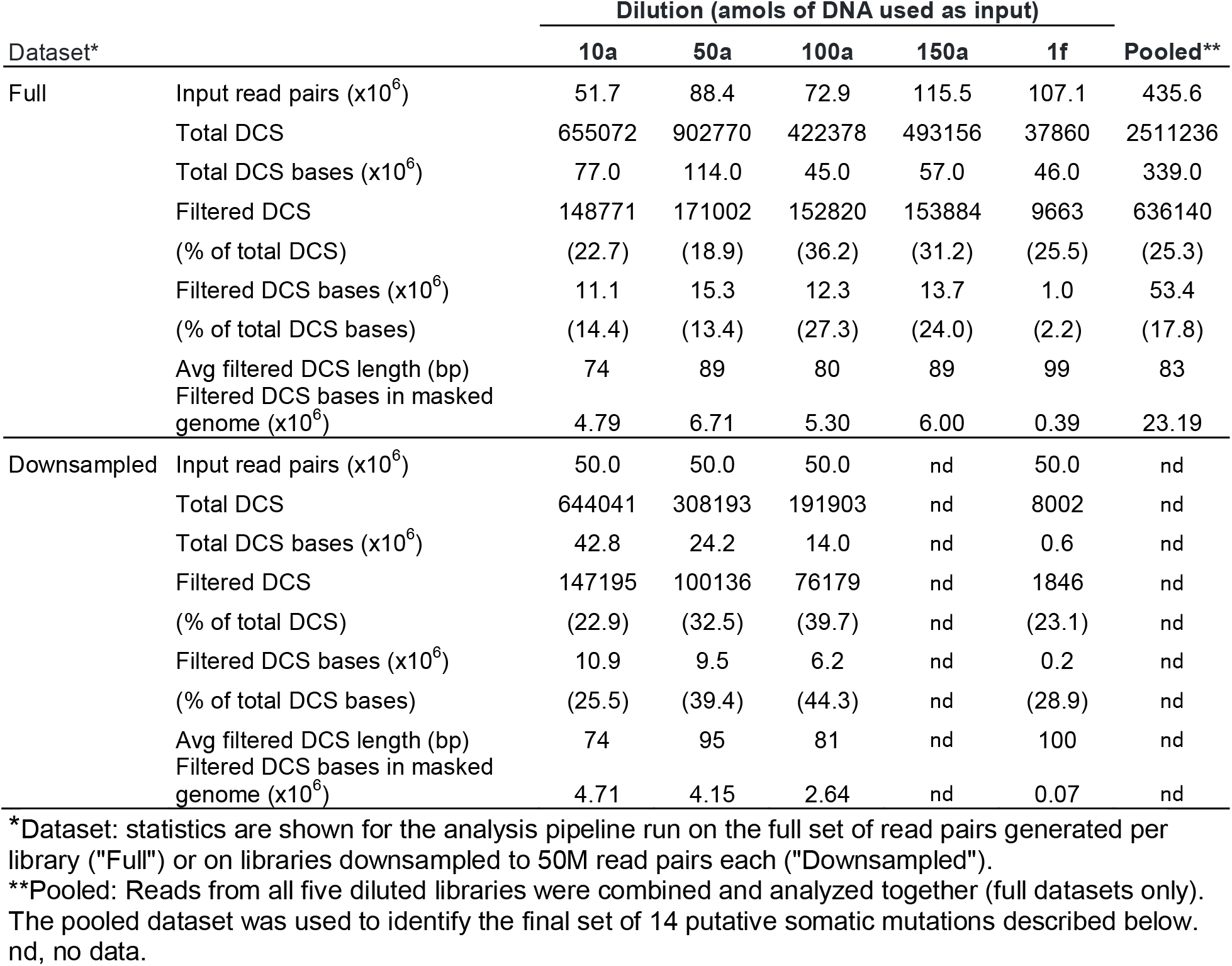
Consensus-making statistics for five dilutions of a *D. magna* gDNA-Seq library.

| Dataset* |  | Dilution (amols of DNA used as input) |  |  |  |  | Pooled** |
| --- | --- | --- | --- | --- | --- | --- | --- |
|  |  | 10a | 50a | 100a | 150a | 1f |  |
| Full | Input read pairs (x10<sup>6</sup>) | 51.7 | 88.4 | 72.9 | 115.5 | 107.1 | 435.6 |
|  | Total DCS | 655072 | 902770 | 422378 | 493156 | 37860 | 2511236 |
|  | Total DCS bases (x10<sup>6</sup>) | 77.0 | 114.0 | 45.0 | 57.0 | 46.0 | 339.0 |
|  | Filtered DCS | 148771 | 171002 | 152820 | 153884 | 9663 | 636140 |
|  | (% of total DCS) | (22.7) | (18.9) | (36.2) | (31.2) | (25.5) | (25.3) |
|  | Filtered DCS bases (x10<sup>6</sup>) | 11.1 | 15.3 | 12.3 | 13.7 | 1.0 | 53.4 |
|  | (% of total DCS bases) | (14.4) | (13.4) | (27.3) | (24.0) | (2.2) | (17.8) |
|  | Avg filtered DCS length (bp) | 74 | 89 | 80 | 89 | 99 | 83 |
|  | Filtered DCS bases in masked genome (x10<sup>6</sup>) | 4.79 | 6.71 | 5.30 | 6.00 | 0.39 | 23.19 |
| Downsampled | Input read pairs (x10<sup>6</sup>) | 50.0 | 50.0 | 50.0 | nd | 50.0 | nd |
|  | Total DCS | 644041 | 308193 | 191903 | nd | 8002 | nd |
|  | Total DCS bases (x10<sup>6</sup>) | 42.8 | 24.2 | 14.0 | nd | 0.6 | nd |
|  | Filtered DCS | 147195 | 100136 | 76179 | nd | 1846 | nd |
|  | (% of total DCS) | (22.9) | (32.5) | (39.7) | nd | (23.1) | nd |
|  | Filtered DCS bases (x10<sup>6</sup>) | 10.9 | 9.5 | 6.2 | nd | 0.2 | nd |
|  | (% of total DCS bases) | (25.5) | (39.4) | (44.3) | nd | (28.9) | nd |
|  | Avg filtered DCS length (bp) | 74 | 95 | 81 | nd | 100 | nd |
|  | Filtered DCS bases in masked genome (x10<sup>6</sup>) | 4.71 | 4.15 | 2.64 | nd | 0.07 | nd |

\*Dataset: statistics are shown for the analysis pipeline run on the full set of read pairs generated per library ("Full") or on libraries downsampled to 50M read pairs each ("Downsampled").
\*\*Pooled: Reads from all five diluted libraries were combined and analyzed together (full datasets only). The pooled dataset was used to identify the final set of 14 putative somatic mutations described below. nd, no data.

| Dataset* | Library parameter | Dilution (amols of DNA used as input) |  |  |  |  | Pooled** |
| --- | --- | --- | --- | --- | --- | --- | --- |
|  |  | 10a | 50a | 100a | 150a | 1f |  |
| Full | Input read pairs ($\times 10^6$) | 51.7 | 88.4 | 72.9 | 115.5 | 107.1 | 435.6 |
|  | Total DCS | 655072 | 902770 | 422378 | 493156 | 37860 | 2511236 |
| | Total DCS bases ($\times 10^6$) | 77.0 | 114.0 | 45.0 | 57.0 | 46.0 | 339.0 |
|  | Filtered DCS | 148771 | 171002 | 152820 | 153884 | 9663 | 636140 |
|  | (% of total DCS) | (22.7) | (18.9) | (36.2) | (31.2) | (25.5) | (25.3) |
| | Filtered DCS bases ($\times 10^6$) | 11.1 | 15.3 | 12.3 | 13.7 | 1.0 | 53.4 |
|  | (% of total DCS bases) | (14.4) | (13.4) | (27.3) | (24.0) | (2.2) | (17.8) |
|  | Avg filtered DCS length (bp) | 74 | 89 | 80 | 89 | 99 | 83 |
| | Filtered DCS bases in masked genome ($\times 10^6$) | 4.79 | 6.71 | 5.30 | 6.00 | 0.39 | 23.19 |
| Downsampled | Input read pairs ($\times 10^6$) | 50.0 | 50.0 | 50.0 | nd | 50.0 | nd |
|  | Total DCS | 644041 | 308193 | 191903 | nd | 8002 | nd |
| | Total DCS bases ($\times 10^6$) | 42.8 | 24.2 | 14.0 | nd | 0.6 | nd |
|  | Filtered DCS | 147195 | 100136 | 76179 | nd | 1846 | nd |
|  | (% of total DCS) | (22.9) | (32.5) | (39.7) | nd | (23.1) | nd |
| | Filtered DCS bases ($\times 10^6$) | 10.9 | 9.5 | 6.2 | nd | 0.2 | nd |
|  | (% of total DCS bases) | (25.5) | (39.4) | (44.3) | nd | (28.9) | nd |
|  | Avg filtered DCS length (bp) | 74 | 95 | 81 | nd | 100 | nd |
| | Filtered DCS bases in masked genome ($\times 10^6$) | 4.71 | 4.15 | 2.64 | nd | 0.07 | nd |

\*Dataset: statistics are shown for the analysis pipeline run on the full set of read pairs generated per library ("Full") or on libraries downsampled to 50M read pairs each ("Downsampled").
\*\*Pooled: Reads from all five diluted libraries were combined and analyzed together (full datasets only). The pooled dataset was used to identify the final set of 14 putative somatic mutations described below. nd, no data.

**Figure 3.**
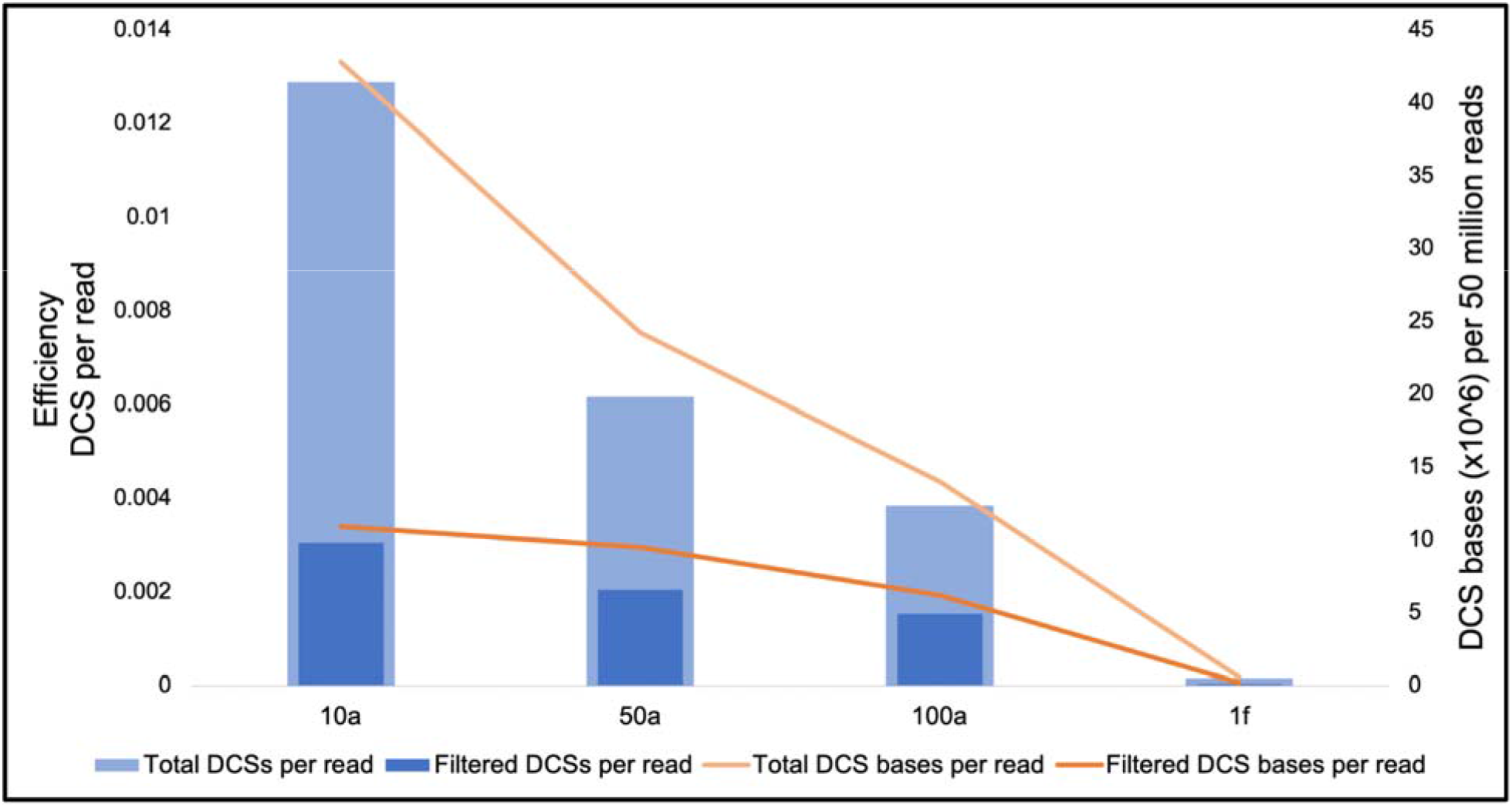
Efficiency of consensus sequence generation by library dilution. Consensus-making efficiency metrics for four diluted libraries of *D. magna* gDNA down-sampled to 50M read pairs per library. On the left, the number of DCSs per input read pair (both before and after DCS filtering steps). Filtering removed DCSs if the underlying read pairs mapped improperly, if the DCS was derived from an insert <20 bp or >500 bp, if the DCS mapped with a low alignment score, or if the DCS mapped equally well to multiple locations in the reference genome [see methods]). On the right, genome coverage based on the 50 M reads is plotted (orange line) for each dilution level.

**Figure 3** shows the relationship between DNA input amount and two different measures of sequencing efficiency (DCSs per read pair and DCS bases per 50 M reads). While efficiency can be assessed based on the total number of DCSs generated per read (total DCSs per read; light blue bars in **Fig. 3**), in practice not all DCSs can be mapped with high confidence (e.g., many map to multiple locations in the genome), and those with ambiguous or irregular mapping must be discarded. Thus DCSs per read after filtering problematic DCSs (filtered DCS per read; dark blue bars in **Fig. 3**) may be a more realistic measure of efficiency, and will vary from project to project due to technical artifacts associated with library construction, reference genome assembly, and mapping. A related measure of efficiency is the DCS bases per 50 million reads, which gives an indication of the breadth of genomic space that can be surveyed with a given amount of sequencing power (orange lines; **Fig. 3**).

The 10a library had the highest raw (unfiltered) DCS making efficiency, generating approximately one DCS per 75 input read pairs (1.34% efficiency). The 50a library was roughly half as efficient (0.63% efficiency, ca. one DCS per 160 read pairs), but still performed better than the 100a and 1f libraries (efficiencies of 0.38% and 0.02%, ca. one DCS per 260 and 5000 read pairs, respectively). The sequencing efficiency we observed at 10 amol is similar to that obtained in *Salmonella* (∼1.8%) by Matsumura et al. [9], and in the range (1-10%) typically obtained by Kennedy et al. [7]. Our efficiencies, however, were consistently lower than those of Matsumura et al. [9] at higher input amounts, and while Matsumura et al. [9] found an optimal dilution range of 39-156 amol for 50M read pairs, our distribution of sequencing efficiencies suggests that the optimal dilution is at or below 10 amol (**Fig. 3**).

The difference between our optimal dilution and that of Matsumura et al. [9] may be explained in part by experimental error in the estimation of DNA concentrations - for example, Matsumura et al. used Bioanalyser-based estimates of library concentration whereas we used Qubit-based estimates. The primary explanation for the observed differences, however, is likely that we required a minimum of two read pairs per SSCS whereas Matsumura et al. [9] only required one. Thus, our DCS assembly strategy was more stringent, but at the expense of fewer total DCSs per input. Our requirement for two read pairs per SSCS was also used by Hoang et al. [8], whereas Kennedy et al. [7] require three read pairs, so our approach represents a middle ground between previously published methods with regard to stringency. Finally, another likely explanation is that our reference genome assembly is less complete and of lower quality than those used by Matsumura et al. [9], resulting in lower mapping efficiency, which is consistent with the fact that there were large differences between total DCS per read and filtered DCS per read (**Fig. 3**). The 10a library also yielded the most DCS bases per read (**Fig. 3**). It is possible that even higher efficiency without a reduction in coverage might have been obtained by using less than 10 amol of input DNA. The order of magnitude between our optimum and that reported in Matsumura et al. [9] highlights the fact that the optimal input will vary depending on both the quality of the reference genome and the consensus-building criteria, and should be determined empirically rather than simply taken from other studies.

### Variants were filtered by masking the genome

We next used the full datasets (not downsampled to 50M read pairs) from all five dilutions (**Table 1, Fig. S1**) to identify variants and estimate a genome-wide somatic mutation rate. Across the five libraries, consensus reads identified 126,670 putative variants (**Fig. 4**), an unrealistically large number. Consequently, we identified several criteria by which to identify and remove likely artifacts (**Fig. 4**). First, the majority of these variants (73%) were removed by excluding known heterozygous sites. Of the remaining 34,245 surviving this filter, many were densely clustered in regions with unusually high coverage in the reference genome assembly. High read coverage and variant frequency is characteristic of assembly errors in reference genomes assembled using short read data, such as ours. Due to the difficulty of accurately constructing contigs of repetitive DNA out of short reads, many repetitive regions are artificially collapsed into a single locus. If multiple copies are subsequently recovered in Duplex Sequencing, they will falsely map to one locus, and the erroneous mapping will cause sequence differences to be called as variants.

**Figure 4.**
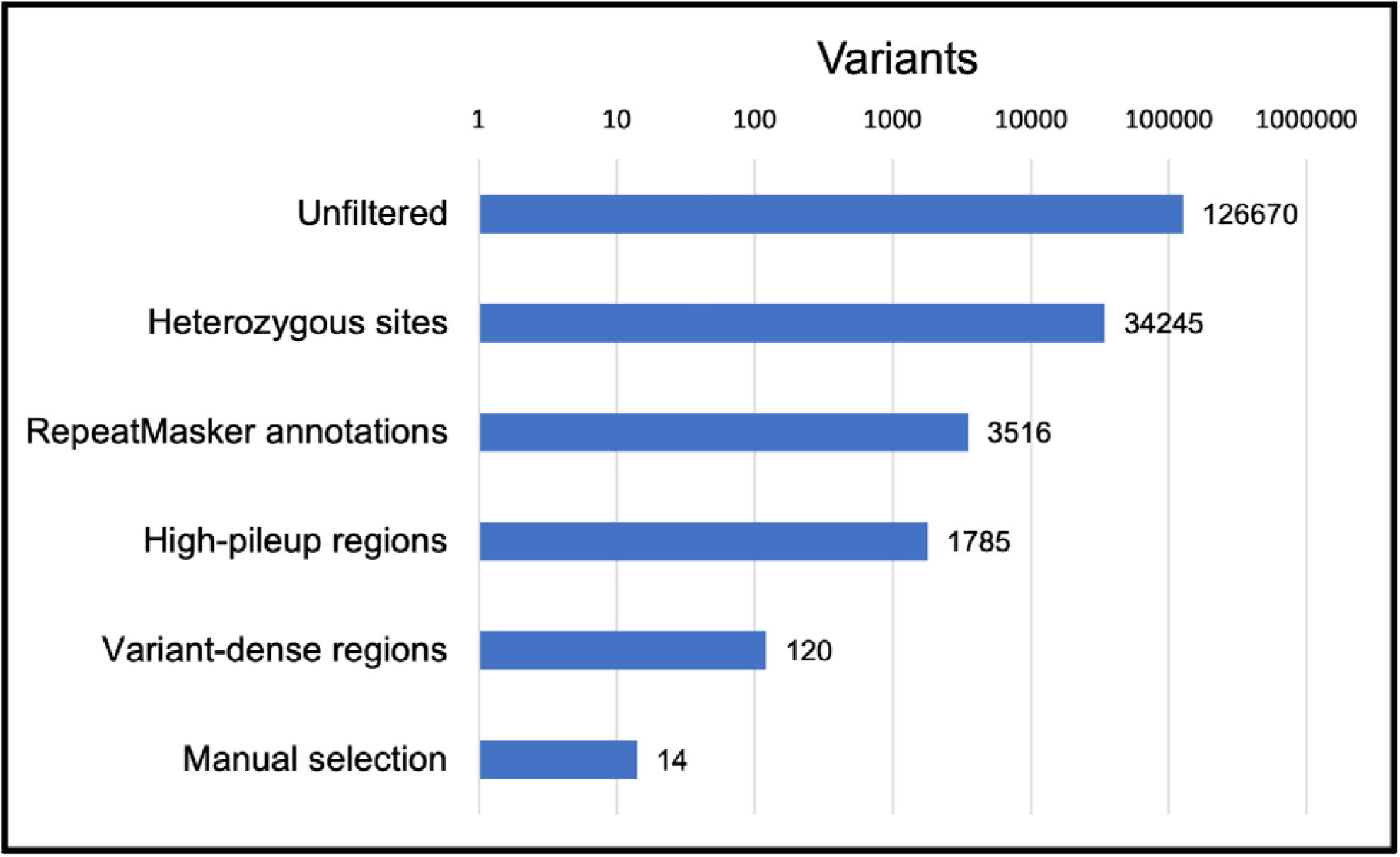
Variants representing putative somatic mutations sequentially excluded by each masking filter. Numbers indicate how many variants remained after removing heterozygous sites, regions annotated by RepeatMasker, windows with high read pileup in the original reference genome, windows with more than 3 variants per 500 bp, and then variants manually excluded.

Fortunately, the presence of such regions in the reference genome can be diagnosed using tools to identify repeats. We applied three filters to mask repetitive regions (**Fig. 4**). First, we masked regions flagged by RepeatMasker and RepeatModeler [22,23]. Then, we masked regions of the original reference assembly with excessive read coverage (more than double the median). Lastly, regions of the genome with more than 3 variants per 500 bp sliding window were masked, as such sites are likely to represent genome assembly and/or mapping errors. After all masking steps, 45.3 Mbp of the 120 Mbp genome assembly remained (38%). Within these 45.3 Mbp, 21 Mbp (47%) were covered by a total of 23.2 Mbp of DCSs from all libraries combined (**Table 1**), meaning we were able to detect somatic mutations in 17.5% of the sequenced genome.

Collectively, these filters removed >99.9% of variants from the initial, unmasked set (**Fig. 4**), leaving 120 putative somatic mutations. These variants were manually inspected in IGV (Thorvaldsdottir et al., 2013) to exclude variants with a variant allele fraction over 0.5 in cases where a locus was covered by multiple DCSs, or where the reads used to build the reference genome showed heterozygosity at that locus. Variants in regions of the reference assembly lacking read coverage (regions where the sequence comes from the genome used as a scaffold [17]) were also excluded. This left a final set of 14 putative somatic variants in the approximately 18% of the genome in which we were able to look for somatic mutations.

### 93% of putative somatic variants were confirmed to not be germline mutations

As noted by Hoang et al. [8], because somatic mutations are both ephemeral and typically found at extremely low VAF, they are, potentially, impossible to validate. However, it is possible to use other methods to ensure that variants detected are not, in fact, germline mutations. We assessed the validity of the final set of 14 putative mutations by Sanger sequencing using the leftover fractions of each library not used for Illumina sequencing. For each mutation, we PCR amplified and Sanger sequenced the relevant region in both the original, undiluted library, as well as in the relevant diluted library from which the mutation was called (10 amol, 50 amol, 100 amol, 150 amol, or 1 fmol). In 13 out of 14 cases (93%), the mutation was not observed in the undiluted library, confirming that the mutation is not a germline variant. In the 14th case, the undiluted library only exhibited the putative mutation, indicating that the putative variant was not a somatic mutation, and that it is either a mis-call in the reference genome or a *de novo* germline mutation that became fixed in the population subsequent to sequencing the reference genome.

None of the 13 non-germline mutations were observed in the Sanger traces from their originating diluted library. Thus, we were not able to demonstrate definitively that any of the called variants are true somatic mutations. This is not surprising given that somatic mutations are often present at very low VAF (<<1%), potentially affecting only one or a few cells. Although the bottlenecking step would be expected to greatly increase the VAF, we estimate that our highest input library (1 fmol) started with an amount of DNA equivalent to roughly 987-1909 genome copies, and our lowest input library (10 amol) started with an amount of DNA equivalent to 9.8-19.9 genome copies (See Supplementary Results, **Table S1**). Therefore, even in the 10 amol library, recovered somatic variants could affect less than 10% of copies at the locus in question, and it is unlikely that such variants would present discernible peaks above background in Sanger traces. Consequently, though we established that 13 out of 14 of the putative mutations are not previously undetected germline variants, the data are inconclusive as to whether they represent false positives in the bottlenecked libraries or true variants at VAFs below the detection limit of Sanger sequencing. However, because Duplex Sequencing requires evidence from both DNA strands to call a mutation, complementary errors affecting both DNA strands, either during PCR or sequencing, would be required for artifacts to be called as variants. The likelihood of such “jackpot” errors is less than one per 1 billion bases sequenced [6], so we can be reasonably confident that these 13 putative variants represent true, *in vivo* mutations (but see Supplementary Information for discussion of potential sources of error associated with endogenous UMIs, and especially library preparation, as well as recently proposed modifications to further lower false positive rates).

### The estimated somatic mutation rate is two orders of magnitude higher than the germline mutation rate

The 13 somatic variants identified in 23.2 Mbp of analyzed consensus sequence yields a mutation rate of 5.6 × 10^−7^ per base pair. This rate is ca. 6-60 times higher than the reported error rate for Hawk-Seq [9], and 560 times higher than the theoretical error rate for Duplex Sequencing [6], suggesting that errors have not inflated the estimate appreciably. Furthermore, our estimate is roughly two orders of magnitude higher than the germline rate of 3.60 × 10^−9^ estimated by Ho et al. [17]. This ratio of somatic to germline mutation rates is similar to those found in both humans and mice [26], lending plausibility to our somatic mutation rate estimate.

By necessity, we sampled only a subset (18%) of the genome, so our mutation rate may differ from the true average rate for the entire genome. Mutation rates are known to vary greatly across genomic features, and repetitive elements such as simple repeats and transposable elements are likely to mutate at a rate that differs from the genomic average due to factors, such as transcription level and chromatin status [27]. Our mutation rate therefore provides an estimate of the frequency of mutations in the less repetitive subset of the *D. magna* genome. Though this rate may differ from that of the whole genome, the sampled fraction is enriched for functional elements (e.g., genes and regulatory sequences) so the rate we obtained is likely to be of the greatest relevance to clinical and basic biology.

## Conclusion

Duplex Sequencing is one of the most sensitive and error-free methods for detecting rare somatic mutations [3,5,7,14]. However, this approach has not previously been applied to organisms lacking high-quality reference genomes. Here, we presented guidelines for optimizing library dilution for efficient Duplex Sequencing, and applied this approach to call rare variants in *D. magna*, a species with a draft reference assembly. Due to the draft quality genome, it was necessary to mask problematic regions of the assembly from analysis in order to separate signal from noise. We masked regions of the reference assembly with repeat annotations, unusually high contributing read coverage, and unusually high variant density, drastically reducing the number of false positives stemming from mapping errors. We detected a per-base somatic mutation rate approximately two orders of magnitude higher than the germline mutation rate for the same genotype of *Daphnia*. Thus, we have added to the short list of species for which genome-wide rates of both germline and somatic mutation rates have been estimated. This ratio of germline to somatic mutation rates is in line with those found in other species, suggesting that SMS, in combination with our strategies for filtering variants, enables accurate estimation of somatic mutation rates in organisms with imperfect genome assemblies.

## Abbreviations used

DCS: Double Strand Consensus Sequence
NGS: Next Generation Sequencing
SMS: Single Molecule Sequencing
SSCS: Single Strand Consensus Sequence
UMI: Unique Molecular Identifier
VAF: Variant Allele Frequency

## Acknowledgements

We thank Brendan F. Kohrn and Scott R. Kennedy for extensive advice and consultation on running the Duplex Sequencing pipeline. We thank Eddie K.H. Ho for advice and assistance on masking the *D. magna* reference genome sequence. We would also like to acknowledge our funding sources: awards from Reed College (to ES) and grants from the National Institute of General Medical Sciences of the National Institutes of Health (GM132861) and National Science Foundation (MCB-1150213) to SS.

## Supplementary Information

**Figure S1**. Consensus-making efficiency (DCS per input read pair) for five diluted libraries of D. magna gDNA without down-sampling. DCS were not filtered for insert size or alignment score.

### Supplemental Results

Estimates of genome copies per library

**Figure S2**. Bioanalyzer trace for the undiluted and unamplified NEBNext Ultra II FS library used as source for the 5 bottlenecked libraries (average fragment size estimated to be 456bp).

**Table S1**. Estimates of the number of genome equivalents per DNA library.

### Supplemental Discussion

Potential sources of error, and modifications to, Duplex Sequencing to further refine somatic mutation rate estimates

