## Supplementary Information for "Estimating somatic mutation rates by Duplex Sequencing in non-model organisms: *Daphnia magna* as a case study"

**Table of Contents**

**Figure S1.** Consensus-making efficiency (DCS per input read pair) for five diluted libraries of D. magna gDNA without down-sampling. DCS were not filtered for insert size or alignment score.	**2**

**Estimates of genome copies per library**	**2**

**Figure S2.** Bioanalyzer trace for the undiluted and unamplified NEBNext Ultra II FS library used as source for the 5 bottlenecked libraries (average fragment size estimated to be 456bp).	**3**

**Table S1.** Estimates of the number of genome equivalents per DNA library.	**3**

**Potential sources of error, and modifications to, Duplex Sequencing to further refine somatic mutation rate estimates**	**4**

**Bioinformatics scripts**	**5**

**Supplementary References**	**7**

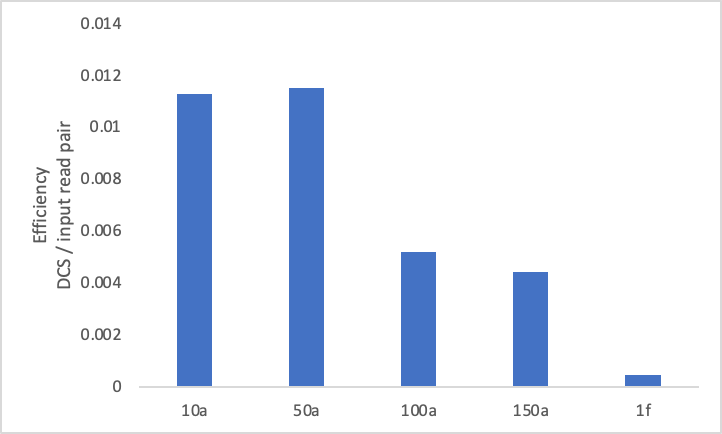

### **Figure S1.** Consensus-making efficiency (DCS per input read pair) for five diluted libraries of *D. magna* gDNA without down-sampling. DCS were not filtered for insert size or alignment score.

### *Estimates of genome copies per library*

The undiluted and unamplified library that was used as input for the five bottlenecked libraries had an average fragment size of 456bp (**Fig. S2**). The number of molecules per diluted library was calculated as the molar mass of input DNA x Avogadro’s number (6.022 x 10^23^). The number of base pairs of DNA per library was then calculated by multiplying the number of molecules by the average length (456bp). Finally, the number of genome equivalents per library was estimated by dividing the total number of base pairs in the library by the number of base pairs in the *D. magna* genome. Two published estimates for the size of the *D. magna* genome vary considerably, so we estimated genome copy number using each - 123 Mbp [[1]](https://www.zotero.org/google-docs/?peB4jk) and 238 Mbp [[2]](https://www.zotero.org/google-docs/?KTaskl). Values are shown in **Table S1**.

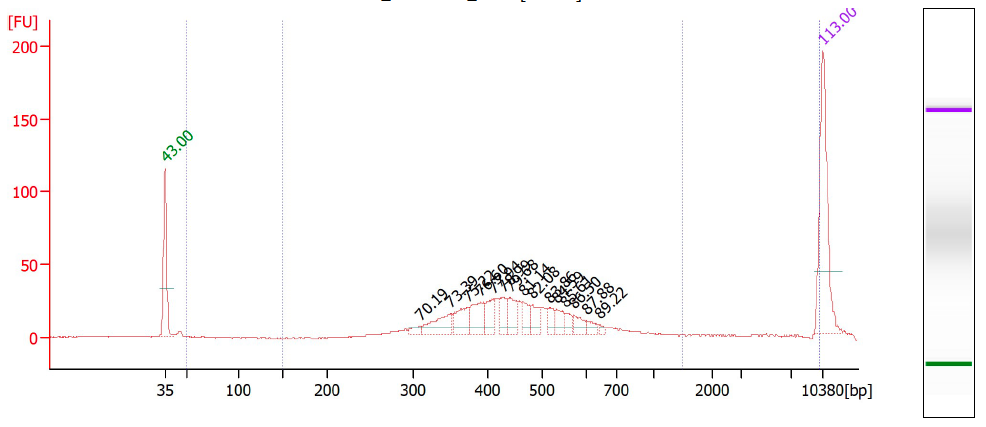

#### **Figure S2.** Bioanalyzer trace for the undiluted and unamplified NEBNext Ultra II FS library used as source for the 5 bottlenecked libraries (average fragment size estimated to be 456bp).

#### **Table S1.** Estimates of the number of genome equivalents per DNA library.

|  |  |  |  | # genome equivalents^3^ | |
| --- | --- | --- | --- | --- | --- |
| Input DNA | Input DNA (moles) | # molecules^1^ | # BPs^2^ | Routtu et al. 2014 | Lee et al. 2019 |
| 1 fmole | 1E-15 | 6.02E+08 | 2.35E+11 | 986.8 | 1909.4 |
| 150 amoles | 1.5E-16 | 9.03E+07 | 3.52E+10 | 148.0 | 286.4 |
| 100 amoles | 1.00E-16 | 6.02E+07 | 2.35E+10 | 98.7 | 190.9 |
| 50 amoles | 5.00E-17 | 3.01E+07 | 1.17E+10 | 49.3 | 95.5 |
| 10 amoles | 1.00E-17 | 6.02E+06 | 2.35E+09 | 9.9 | 19.1 |

^1^# molecules = moles * 6.022E23

^2^# BPs = # molecules * 390bp/molecule (average fragment size = 456bp minus 66bp of adapater, leaving 390bp of genomic DNA)

^3^# genome equivalents = # BPs / genome size (bp) [Values are given for a genome size estimate from flow cytometry (238 Mbp; [[2]](https://www.zotero.org/google-docs/?psSx1U)), and from a recently published genome sequence (123 Mbp; [[1]](https://www.zotero.org/google-docs/?c3NIuj))

### *Potential sources of error, and modifications to, Duplex Sequencing to further refine somatic mutation rate estimates*

By randomly surveying the entire genome in a mostly unbiased fashion, and not restricting sequencing to small target regions, bottleneck sequencing has the advantage of not requiring exogenous UMIs. When randomly surveying a typical eukaryotic genome, it is extremely unlikely that two sampled DNA fragments will have sheared at the exact same genomic coordinates. Consequently, one can use the end sequences and mapping coordinates of reads as molecular identifiers to group read families (endogenous barcoding), rather than ligating UMI adapters to DNA fragments prior to sequencing (exogenous barcoding). By enabling the use of endogenous UMIs, bottlenecked Duplex Sequencing streamlines library preparation. However, the likelihood that two copies of a given DNA molecule will have the same breakpoints (“overlap by accident” or “OBA”; [[3]](https://www.zotero.org/google-docs/?rlI5Fr)) increases with decreasing amounts of DNA examined (i.e, when performing target enrichment or in organisms such as bacteria with small genome sizes). OBA is also expected to increase with the use of fragmentation methods that do not shear DNA randomly (e.g., sequence-specific restriction enzymes such as those recommended by Abascal et al. [[4]](https://www.zotero.org/google-docs/?KSgdQ8)). OBA can cause genuine mutations to be filtered out as artifacts, thereby reducing estimates of mutation rate [[3]](https://www.zotero.org/google-docs/?u5VxpG). Thus, exogenous UMIs should be considered when working with small genomes, targeted regions of larger genomes, or when utilizing non-random fragmentation methods.

Though the theoretical error rate for Duplex Sequencing is <10^-9^/bp [[5]](https://www.zotero.org/google-docs/?GE4mGx), the rate has been shown to be higher in practice (10^-4^ to 10^-7^; [[6]](https://www.zotero.org/google-docs/?3XmH5W). The higher than expected error rate is thought to stem largely from DNA damage induced during the DNA fragmentation step of standard Duplex Sequencing library preparation [[6]](https://www.zotero.org/google-docs/?izwTci). Specifically, fragmentation generates single strand overhangs, as well as internal nicks and gaps, and where these single stranded regions are damaged, the end repair process converts them to double-stranded errors [[4,6]](https://www.zotero.org/google-docs/?RsiuP9). Abascal et al. [[4]](https://www.zotero.org/google-docs/?fBRGx7) estimated that sonication and subsequent end repair introduced ca. 1200 mutations per diploid human cell, significantly inflating estimates of mutation rate by BotSeqS [[7]](https://www.zotero.org/google-docs/?XS3d6o).

Because these artifacts are primarily the result of end-repair, removing several base pairs from the ends of consensus reads has been shown to considerably reduce the number of artifacts. For example, You et al. [[6]](https://www.zotero.org/google-docs/?V2g6Vt) showed that the 7 terminal base pairs on each end of a consensus read have elevated error rates, and removing these bases cut the resulting estimate of mutation rate in half. We took a more conservative approach and removed the first and last 15 bp from each DCS. Thus, our data should be largely free of terminal end repair artifacts. Alternative approaches include the use of single strand-specific nucleases to blunt 3’ overhangs [[8]](https://www.zotero.org/google-docs/?2iEEBF) or using blunt-end restriction enzyme-based fragmentation [[4]](https://www.zotero.org/google-docs/?TsIl4H), eliminating the need for end repair.

However, fragmentation also produces internal nicks and gaps, so eliminating or computationally removing 3’ ends of consensus sequences does not remove all artifacts [[4,6]](https://www.zotero.org/google-docs/?yu2Hui). To some extent, our approach may have reduced the occurrence of these errors because we utilized enzymatic fragmentation which is less damaging than mechanical methods [[6]](https://www.zotero.org/google-docs/?GB4iAj), but enzymatic fragmentation is nonetheless known to cause nicks and gaps [[9]](https://www.zotero.org/google-docs/?8a8KlT). Thus, additional strategies aimed at mitigating the errors associated with this type of damage should reduce error rates and improve overall estimates of somatic mutation rate. Abascal et al. [[4]](https://www.zotero.org/google-docs/?RDxp15) described a modified form of Duplex Sequencing, - Nanorate Sequencing or NanoSeq - which, in addition to generating blunt ended DNA fragments, also includes non-A dideoxynucleotides (ddBTPs) in the A-tailing reaction. As a consequence, most nick extension reactions will produce non-amplifiable fragments that will not be sequenced. Abascal et al. [[4]](https://www.zotero.org/google-docs/?n3gAEb) estimate that NanoSeq achieves an error rate less than 5 x 10^-9^/bp, approaching the theoretical error rate of Duplex Sequencing [[5]](https://www.zotero.org/google-docs/?ejVL62). Thus, although restriction enzyme-based fragmentation does not give random sampling of the entire genome, NanoSeq is likely to yield more accurate estimates of the somatic mutation rate than other SMS approaches.

### *Bioinformatics scripts*

Pipeline and dependencies setup

a. #Install miniconda
   i. $ wget <https://repo.anaconda.com/miniconda/Miniconda3-latest-Linux-x86_64.sh>
   ii. $ bash Miniconda3-latest-Linux-x86_64.sh
b. Install mamba via conda install
   i. $ conda install -y -c conda-forge mamba
c. #Install all other dependencies via mamba install
   i. $ mamba install -y -c bioconda -c conda-forge snakemake=5.25.0
   ii. $ mamba install -y -c bioconda -c conda-forge pandas
   iii. $ mamba install -y -c bioconda bwa
   iv. $ mamba install -y -c bioconda -c conda-forge blast=2.6.0
   v. $ mamba install -y -c bioconda -c conda-forge samtools
   vi. $ mamba install -y -c bioconda -c conda-forge picard
d. #Download pipeline and update to latest version
   i. $ git clone <https://github.com/KennedyLabUW/Duplex-Seq-Pipeline.git>
   ii. $ git pull
   iii. $ git checkout -f v2.0.0_prerelease
   iv. $ rm .env_initialized
   v. $ bash setupDS.sh [#cores]

Reference genome and bed file setup

e. #Index reference genome
   i. $ bwa index IASC_sorted.fasta
   ii. $ samtools faidx IASC_sorted.fasta
   iii. $ picard CreateSequenceDictionary R=IASC_noNs.fasta O=IASC_noNs.dict
f. #Create and format bed file covering entire reference genome
   i. $ bioawk -c fastx '{print $name"\t0\t"length($seq)}' input.fasta > output.bed
   ii. $ awk '{print $0,"name"NR,".","."}' output.bed > output_6col.bed
   iii. $ tr -s " " "\t" < output_6col.bed > genome.bed

Run pipeline

g. #Run pipeline. Navigate to directory below the main pipeline directory but above the directory containing the input files and run the pipeline. Config.csv should be edited to the desired parameters of the run.
   i. $ bash ../DS config.csv
a. #Call variants
   i. $cd /path/to/run_directory/Final/dcs
   ii. $ for i in *dcs.final.bam; do bcftools mpileup -f /vol_b/RefGenome/IASC_sorted.fasta $i | bcftools call -mv -V indels Ov -o $i.vcf; done
b. #Generate mutational spectra
   i. $ python helmsman.py --input /path/to/.vcf --fastafile /vol_b/RefGenome/IASC_sorted.fasta --projectdir /vol_b/Figures/vcfs/snps_vcf/helmsman_out

Filter variants

\# Exclude known heterozygous sites

\# Find and exclude intersection of initially called variants and known heterozygous sites

$ bedtools intersect -u -a XS_i_0.vcf.bed -b IA.BISNP.vcf.bed

\# Exclude regions obtained by running RepeatMasker

\# Extend windows excluded by RepeatMasker by 500 bp

$ bedtools slop -b 500 -i IASC.5000.fasta.out.bed -g /vol_b/RefGenome/IASC_sorted.genome > RepeatMasker_exlusions_slop500.bed

\# Exclude regions where original reads used to build reference genome have over 2x expected coverage in reference genome.

\# map reads used to build reference assembly back to reference

$ bwa mem -t 24 /vol_b/RefGenome/IASC_sorted.fasta IASC.R1.fastq IASC.R2.fastq > IASC_remapped.sam

$ samtools view -S -b IASC_remapped_fixed.sam | samtools sort -o IASC_original_mapped.bam -

\# create coverage graph and get intervals with unusually high coverage

$ bedtools genomecov -ibam IASC_original_mapped.bam -bg > IASC_original_remapped_bg.bed

$ awk '$4 > 80' IASC_original_remapped_bg.bed > coverage_over80_IASC_original.bed

$ bedtools slop -b 500 -i coverage_over80_IASC_original.bed -g /vol_b/RefGenome/IASC_sorted.genome | bedtools sort -i stdin | bedtools merge -i stdin > coverage_over80_IASC_slop500.bed

\# Exclude 500bp windows containing more than one variant

\# divide genome into 500 base pair windows

$ bedtools makewindows -w 500 -s 100 -b IASC_sorted.bed > IASC_sorted_windows_w500_s100.bed

\# calculate variants per window and get windows with more than one variant per window

$ bedtools coverage -a /vol_b/RefGenome/IASC_sorted_windows_w500_s100.bed -b bwa_mem_all_libraries_consensuses.vcf.bed > windows_coverage_IASC_sorted_all_libraries_vcfs.bed

$ for i in *.vcf.bed; do bedtools coverage -a /vol_b/RefGenome/IASC_sorted_windows_w500_s100.bed -b $i | awk -F $'\t' '$4 > 1' > $i.dense_variants.bed; done

$ bedtools slop -b 1000 -i bwa_mem_all_libraries_consensuses.vcf.bed.dense_variants.bed -g /vol_b/RefGenome/IASC_sorted.genome > bwa_mem_all_libraries_consensuses.vcf.bed.dense_variants_slop1kb.bed

\# Merge filters and filter mutations

$ cat * | cut -f -3 | bedtools sort -i stdin | bedtools merge -i stdin > beds_to_exclude_1.38.45.5.bed

$ bedtools subtract -a /vol_b/RefGenome/IASC_sorted.bed -b beds_to_exclude_1.38.45.5.bed > IASC_sorted_excluding_1.38.45.5.bed

$ bedtools intersect -u -a /vol_b/eddie/bwa_mem_all_libraries_consensuses.vcf.bed -b /vol_b/eddie/beds_to_be_merged/temp2/IASC_sorted_excluding_1.38.45.5.bed > all_libraries_vcfs_excluding_1.38.45.5.bed

List of dependencies:

- Miniconda (latest version)
- Mamba
- Python 3.6+
- Snakemake 5.25.*
- Pandas
- bwa 0.7.17.*
- ncbi blast 2.6.0 (only if doing decontamination)
- Samtools
- Picard
- Java
- bcftools
- bedtools
